# Breeding of a high-yield strain for commercial cultivation by crossing *Pholiota adiposa* and *P. limonella*

**DOI:** 10.1101/728626

**Authors:** Chengbo Rong, Shuang Song, Li Yang, Jiachan Zhang, Yurong Niu, Xuejiao Pan, Qinggang Song, Yu Liu, Shouxian Wang

**Author notes:** Chengbo Rong and Shuang Song have contributed equally to this work. Co-corresponding author: E-mail address (Yu Liu). Co-corresponding author: E-mail address (Shouxian Wang).

## Abstract

*Pholiota adiposa* is an edible mushroom with excellent nutritional and medicinal properties. However, fruiting body yields are low, and the commercial cultivation potential of this fungus is limited. In the present study, 279 crossbred strains were obtained by mono-mono crossing of monokaryotic strains derived from *P. adiposa* HS5 and *P. limonella* HS4. Ligninolytic enzymes and mycelial growth rate were used as markers to screen the crossbred strains, and 18 were selected for further analysis. Crossbred strain A10B4 displayed the highest yield, i.e., 165.91 ± 12.56 g per bag, which was 31.34 g and 74.48 g more than that of strains HS5 and HS4, respectively. The mycelial colonization time of A10B4 was 25.18 ± 1.33 days, which was 5.64 days shorter than that of HS5. A10B4 was characterized by inter-simple sequence repeat molecular markers and antagonism tests. Differences in PCR products from parental and crossbred strains were observed. Therefore, the newly developed hybrid strain A10B4, named *P. adiposa-limonella* HS54 and having a high yield and desirable traits, might be suitable for commercial cultivation.

## Introduction

*Pholiota adiposa* (Batsch) P. Kumm. is an edible mushroom that is widely distributed on dead poplars, willows, or birches in forested areas in China [1, 2]. It is also a lignin-degrading macrofungus with excellent nutritional and medicinal properties [2]. Compounds extracted from the fruiting bodies or mycelia of *P. adiposa* display a variety of important biological activities, such as antitumor [2–4], antioxidative [5], antimicrobial [6], and anti-HIV-1 [7] effects. *P. adiposa* has become popular not only in China but also in several regions of Asia, Europe, and North America, owing to its delicious taste and beneficial properties [7, 8].

Mushroom breeding involves various methods, but crossbreeding is considered the most efficient way to develop new, good quality strains of edible mushrooms [9–11]. Intra-species crossbreeding has been reported in *Sparassis latifolia*, *Pleurotus tuoliensis*, *P. eryngii* species etc [10–13], and inter-species crossbreeding has been reported between *P. eryngii* and *P. nebrodensis* etc [14]. High yield is a key benefit of crossbreeding. Most *P. adiposa* cultivars were domesticated from the wild and their yields are not very high, and reports on breeding of this mushroom are limited. In our previous work, strain *P. adiposa* HS5 (JZB2116005) [1], which displayed the highest biological efficiency (BE, 67.88 ± 1.33%, with fruiting bodies in two flushes) and mycelial growth rate (MGR, 2.56 ± 0.03mm/d), was screened as one of five wild strains in comparative studies. *P. limonella* (Peck) Sacc. strain HS4 demonstrated an approximately 1.5-fold higher MGR than HS5 and was closely related to *P. adiposa*. Therefore, to obtain a *P. adiposa* strain with high yield and MGR, we performed inter-species crossbreeding of these two strains.

In *Pholiota* breeding to date, no effective and rapid marker-assisted selection strategies, except for clamp connection, have been developed to evaluate crossbred strains with important agronomic traits at the mycelial growth stage. Laccase is a ligninolytic enzyme. Many edible fungi such as *Volvariella volvacea* [15], *Lentinula edodes*[16, 17], and *Pleurotus florida* [18] have been shown to secrete laccase. Laccase has been shown important to decolorization of Remazol Brilliant Blue R (RBBR) and to lignin degradation and fruiting-body formation [19]. The close relationships between laccase activity, induced primordium differentiation, and development of fruiting bodies were reported by Yang et al [20]. Sun and Xu used decolorization of RBBR by a ligninolytic enzyme as a marker to screen protoplast fusants, and the growth rates and biological efficiencies of new strains were higher than those of the parental strains [19, 21]. Therefore, RBBR offers an efficient approach to evaluating new fungal strains.

In this study, crossbreeding between *P. adiposa* and *P. limonella* was used to develop a new strain (*P. adiposa-limonella*) with high BE and MGR, with decolorization of RBBR by a ligninolytic enzyme used as a marker to screen crossbred strains.

## Materials and methods

### Strains and growth conditions

The fruiting bodies of *P. adiposa* HS5 and *P. limonella* HS4 were collected from Haidian and Changping districts, respectively, in Beijing, China, and then isolated by tissue culture. The strains were cultured and maintained on potato dextrose broth at 25°C. When required, 1.5% (wt/vol) agar was added to the appropriate medium.

### Single-spore isolation

The method was performed as previously described by Wang et al [13]. with minor modifications. Briefly, fruiting bodies were placed in Erlenmeyer flasks containing 100 ml of sterilized water to generate spore suspensions. One hundred microliters of spore suspension (about 1 × 10^3^ spores/ml) was spread onto potato dextrose agar (PDA) plates and incubated at 25°C for five days for spore germination. Monokaryons originating from monospores were identified based on the absence of clamp connections on mycelia ascertained by microscopy (400× magnification).

### PDA-RBBR plate screening and MGR assessment of monokaryons

The strains were transferred from PDA to PDA-RBBR (PDA medium supplemented with RBBR to the terminal concentration of 0.05 % (w/v) to screen for monokaryons. The degree of RBBR fading was designated 1, 2, 3, or 4. A higher number represented greater RBBR decolorization with a more active ligninolytic enzyme. The MGR was determined by observing the radial growth length of mycelia on PDA-RBBR plates.

### Pairings between single-spore isolates

Inter-strain pairing experiments were performed using random single-spore isolates (SSIs) derived from strains *P. adiposa* HS5 and *P. limonella* HS4. Mating was conducted by placing mycelial blocks opposite a monokaryotic mycelium on PDA. Mycelium fragments were taken from the contact zone between the paired colonies and individually transferred to PDA plates for further incubation. When the colonies grew to 1.0-1.5 cm in radius, the mycelia were monitored using a microscope to identify dikaryotic hybrids by clamp connections.

### Antagonistic activity tests

Antagonistic activity assays were performed as previously described, with minor modifications (Xiang et al., 2016). Briefly, two parental strains and the putative hybrids were co-cultured at 25°C in 9-cm PDA plates, with every two mycelial fragments placed 2 cm apart. Somatic incompatibility reactions were observed after incubation for 10 days.

### Fruiting and cultivation methods

Substrates were prepared with 67% (w/w) water content containing 60% cottonseed hull, 18% sawdust, 15% wheat bran, 5% corn flour, 1% gypsum, and 1% lime and placed in polypropylene bags (17 cm × 33 cm × 0.04 cm) at a packing density of 1,000 g of substrate per bag. The bags were autoclaved at 121°C for 120 min. The inoculated bags were then maintained in a spawn running room at 23-25°C and 50-60 relative humidity under dark conditions. After a complete spawn run, mycelial differentiation was induced by stimulation at 0-5°C for 3-5 days. The bags were then transferred to a fruiting chamber that was maintained at 18 ± 2°C and a relative humidity of 80-90% with 12 h of illumination (300-600 Lux) for 15-20 days to obtain fruiting bodies. Each strain was subjected to three treatments, with 12 bags per treatment. Fruiting bodies in first flushes were harvested, yield was the weight of fresh mushrooms per bag, and the biological efficiency (BE) of each fungal strain was calculated based on the following formula: BE (%) = weight of fresh mushrooms harvested per bag/dry weight of cultivation substrates per bag × 100. Mycelial colonization time was calculated as number of days from inoculation to complete colonization of the substrate by the mycelium in the culture bag.

### Inter-simple sequence repeat analysis

Inter-simple sequence repeat (ISSR) amplification was performed in 20-μl reaction volumes containing 0.5 U of Taq DNA polymerase, 2.5 mmol/l MgCl_2_, 0.2 mmol/l dNTPs, 0.75 μmol/l primer, and 50 ng template DNA. Amplification conditions were as follows: 94°C for 5 min; 35 cycles of 94°C for 30 s, 45°C for 45 s, and 72°C for 90 s; and a final extension for 7 min at 72°C. Primer sequences were as follows: P11, AGAGAGAGAGAGAGAGG, P856, ACACACACA CACACACYA, P23,GAGAGAGAGAGAGAGACT, P857, ACACACACAC ACACACYG

### Statistical analysis

All data were statistically analyzed using SPSS PASW Statistics software version 18. All data were obtained in triplicate, and differences were determined by Duncan’s multiple range testing. Statistical differences were considered significant at the 5.0% level (*P* < 0.05).

## Results

### Selection of *P. adiposa* HS5 and *P. limonella* HS4 monokaryons

One hundred and twenty-six and 103 monokaryons were isolated from *P. adiposa* HS5 and *P. limonella* HS4, respectively. Monokaryons were selected for further study based on analysis of mycelial growth rates and their abilities to decolorization of RBBR (Fig 1). For *P. limonella* HS4, the average MGR of 103 monokaryons was 1.77 mm/d in the first round of screening (data not shown), 21 monokaryons with a fading degree ≥ 3 and MGR ≥ 1.77 mm/d were selected during the second round of screening (Table 1). For *P. adiposa* HS5, the average MGR of 126 monokaryons was 1.66 mm/d in the first round of screening (data not shown), 22 monokaryons with a fading degree ≥ 3 and MGR ≥ 1.66 mm/d were selected during the second round of screening (Table 2).

**Figure 1.**
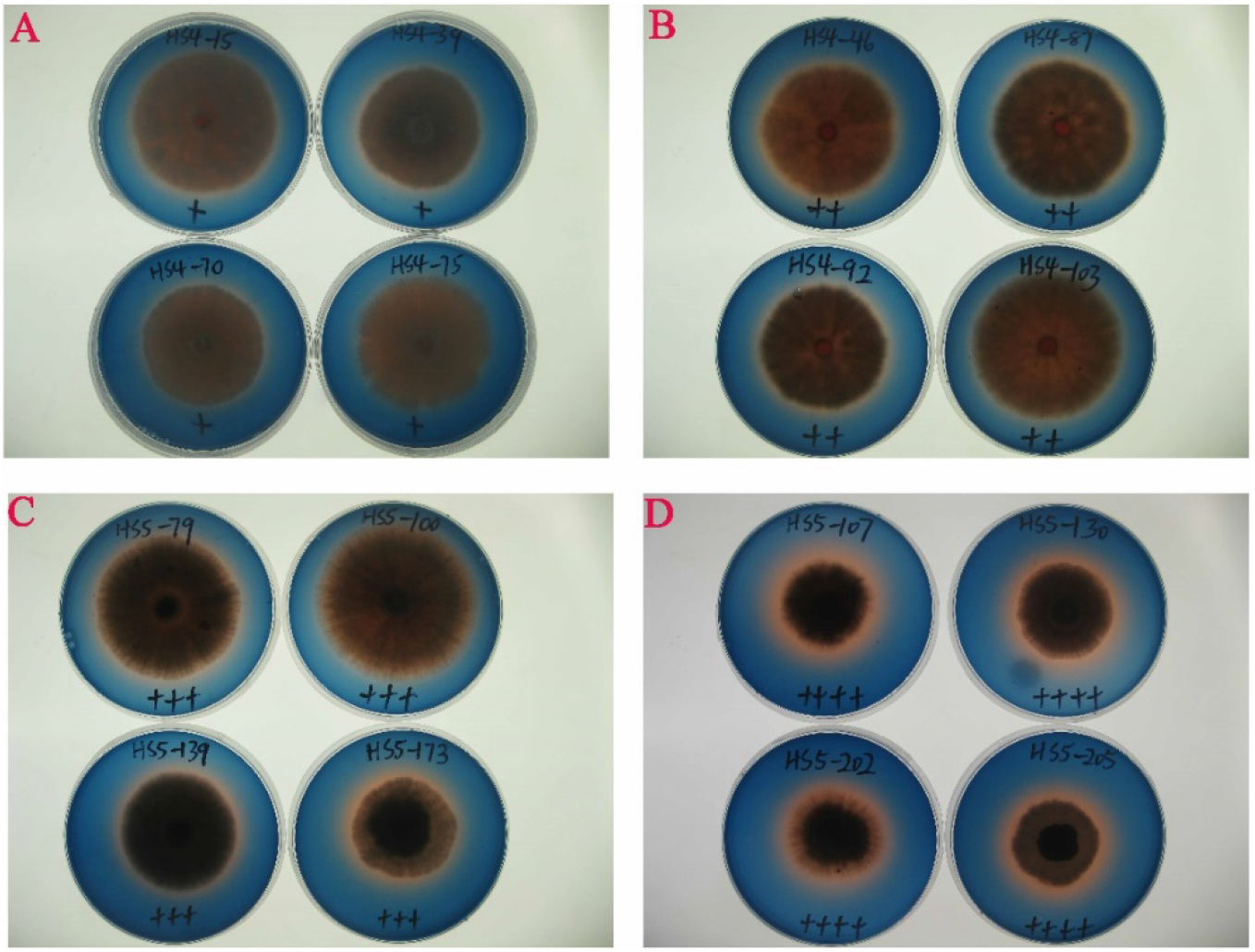
Decoloriztion of HS4 and HS5 monokaryons on PDA plates containing RBBR. The RBBR fading degree was scored as 1 (+), 2 (++), 3 (+++), or 4 (++++). A, B, C, and D show RBBR fading degrees of 1, 2, 3, and 4, respectively.

**Table 1.** Mycelial growth rate and fading degree of *P. limonella* HS4 monokaryons.

| Numbers | Monokaryon number | Mycelial growth rate (mm/d) | Fading degree |
| --- | --- | --- | --- |
| 1 | 45 | 3.10±0.10 | 1 |
| 2 | 65 | 2.76±0.02 | 1 |
| **3** | **82** | **2.70±0.20** | **3** |
| 4 | 103 | 2.53±0.02 | 2 |
| 5 | 15 | 2.52±0.06 | 1 |
| 6 | 75 | 2.49±0.03 | 1 |
| 7 | 87 | 2.48±0.06 | 2 |
| **8** | **72** | **2.42±0.07** | **3** |
| **9** | **90** | **2.42±0.07** | **3** |
| 10 | 78 | 2.39±1.11 | 2 |
| **11** | **43** | **2.39±0.05** | **3** |
| **12** | **100** | **2.39±0.02** | **3** |
| **13** | **42** | **2.38±0.02** | **4** |
| **14** | **101** | **2.33±0.05** | **3** |
| 15 | 46 | 2.26±0.05 | 2 |
| **16** | **107** | **2.24±0.02** | **4** |
| 17 | 92 | 2.19±0.03 | 2 |
| 18 | 39 | 2.18±0.04 | 1 |
| **19** | **9** | **2.17±0.02** | **3** |
| 20 | 70 | 2.16±0.06 | 1 |
| 21 | 28 | 2.16±0.04 | 2 |
| 22 | 32 | 2.14±0.06 | 2 |
| 23 | 10 | 2.13±0.02 | 2 |
| 24 | 30 | 2.13±0.02 | 2 |
| 25 | 16 | 2.12±0.03 | 2 |
| 26 | 40 | 2.11±0.02 | 2 |
| **27** | **102** | **2.11±0.02** | **3** |
| **28** | **84** | **2.10±0.05** | **4** |
| **29** | **50** | **2.10±0.17** | **4** |
| **30** | **62** | **2.05±0.03** | **4** |
| **31** | **95** | **2.05±0.02** | **3** |
| **32** | **41** | **1.99±0.13** | **3** |
| **33** | **76** | **1.98±0.06** | **3** |
| 34 | 29 | 1.96±0.04 | 2 |
| **35** | **51** | **1.93±0.69** | **3** |
| **36** | **67** | **1.89±0.03** | **4** |
| **37** | **68** | **1.88±0.03** | **3** |
| **38** | **2** | **1.85±0.05** | **4** |
| **39** | **19** | **1.77±0.03** | **4** |
| 40 | 52 | 1.72±0.07 | 4 |
| 41 | 83 | 1.63±0.05 | 4 |
| 42 | 25 | 1.56±0.04 | 4 |
| 43 | 5 | 1.51±0.05 | 4 |
| 44 | 3 | 1.47±0.09 | 4 |
| 45 | 4 | 1.44±0.02 | 4 |
| 46 | 89 | 1.44±0.02 | 3 |
| 47 | 71 | 1.36±0.05 | 4 |

Bold letters represented the monokaryons selected for hybridization.

**Table 2.** Mycelial growth rate and fading degree of *P. adiposa* HS5 monokaryons.

| Numbers | Monokaryon number | Mycelial growth rate (mm/d) | Fading degree |
| --- | --- | --- | --- |
| **1** | **100** | **2.77±0.06** | **3** |
| **2** | **61** | **2.51±0.14** | **3** |
| **3** | **60** | **2.47±0.06** | **4** |
| **4** | **79** | **2.38±0.16** | **3** |
| 5 | 3 | 2.31±0.06 | 2 |
| 6 | 43 | 2.28±0.07 | 2 |
| **7** | **76** | **2.19±0.11** | **4** |
| **8** | **17** | **2.09±0.14** | **3** |
| **9** | **139** | **2.07±0.05** | **3** |
| 10 | 32 | 2.01±0.06 | 2 |
| **11** | **29** | **1.97±0.11** | **4** |
| 12 | 11 | 1.95± 0.06 | 2 |
| **13** | **112** | **1.95±0.10** | **3** |
| 14 | 23 | 1.88±0.02 | 2 |
| **15** | **173** | **1.88±0.12** | **3** |
| **16** | **143** | **1.84±0.04** | **4** |
| **17** | **126** | **1.82±0.10** | **3** |
| 18 | 80 | 1.80±0.04 | 2 |
| **19** | **2** | **1.78±0.10** | **4** |
| 20 | 113 | 1.78±0.10 | 2 |
| 21 | 159 | 1.77±0.06 | 1 |
| **22** | **49** | **1.75±0.08** | **4** |
| **23** | **78** | **1.75±0.15** | **3** |
| **24** | **142** | **1.73±0.07** | **4** |
| **25** | **101** | **1.73±0.06** | **4** |
| **26** | **136** | **1.72±0.13** | **3** |
| **27** | **130** | **1.69±0.05** | **4** |
| **28** | **7** | **1.68±0.09** | **4** |
| **29** | **107** | **1.66±0.12** | **4** |
| **30** | **10** | **1.66±0.06** | **4** |
| 31 | 172 | 1.65±0.05 | 1 |
| 32 | 53 | 1.64±0.03 | 4 |
| 33 | 58 | 1.63±0.08 | 4 |
| 34 | 6 | 1.22±0.02 | 4 |
| 35 | 119 | 1.20±0.08 | 2 |
| 36 | 24 | 1.02± 0.30 | 2 |

Bold letters represented the monokaryons selected for hybridization.

### Screening of crossbred strains

Crossbreeding was conducted by mating 22 monokaryons isolated from *P. adiposa* HS5 with 21 monokaryons isolated from *P. limonella* HS4, resulting in a total of 462 matings. Crossbred strains were screened for clamp connections within the contact zone under a microscope. The presence of clamp connections indicates the formation of a dikaryon [12]. A primary screening of these 279 putative crossbred strains was performed on RBBR-PDA (fading degree ≥ 3 and an MGR greater than that of HS5) and resulted in the identification of 55 crossbred strains for further study. A second screening of the 61 strains resulted in 12 putative crossbred strains with fading degree ≥ 3 and an MGR greater than that of HS5 (Table 3). To avoid losing strains with good traits, the top six strains with a high MGR and fading degree ≤ 2 were also selected for further study. All 18 putative hybrids showed strong somatic incompatibility reactions with both parental strains, so these 18 crossbred strains were retained for further study.

**Table 3.** Mycelial growth rate and fading degree of crossbred and parental strains.

| Numbers | Crossbred strains | Mycelial growth rate (mm/d) | Fading degree |
| --- | --- | --- | --- |
| Parental strain | HS4 | 3.46±0.10 | 1 |
| Parental strain | HS5 | 2.29±0.17 | 2 |
| **1** | **A5B1** | **3.14±0.09** | **1** |
| **2** | **A1B18** | **2.81±0.16** | **2** |
| **3** | **A1B17** | **2.66±0.23** | **2** |
| **4** | **A1B11** | **2.57±0.12** | **2** |
| **5** | **A1B12** | **2.57±0.18** | **2** |
| **6** | **A1B6** | **2.53±0.12** | **2** |
| **7** | **A10B5** | **2.47±0.05** | **3** |
| **8** | **A16B3** | **2.46±0.05** | **3** |
| **9** | **A20B3** | **2.44±0.08** | **3** |
| 10 | A13B4 | 2.43±0.03 | 2 |
| **11** | **A8B6** | **2.40±0.12** | **3** |
| 12 | A17B21 | 2.38±0.06 | 2 |
| 13 | A21B6 | 2.38±0.06 | 2 |
| **14** | **A4B6** | **2.36±0.12** | **3** |
| **15** | **A14B4** | **2.36±0.15** | **3** |
| **16** | **A10B4** | **2.35±0.03** | **3** |
| **17** | **A20B12** | **2.35±0.03** | **3** |
| 18 | A8B9 | 2.34±0.08 | 2 |
| **19** | **A6B6** | **2.34±0.04** | **3** |
| 20 | A13B6 | 2.31±0.08 | 2 |
| **21** | **A7B6** | **2.31±0.03** | **3** |
| **22** | **A10B12** | **2.30±0.08** | **4** |
| **23** | **A16B6** | **2.30±0.05** | **4** |
| 24 | A11B2 | 2.29±0.06 | 3 |
| 25 | A14B21 | 2.29±0.04 | 3 |
| 26 | A4B5 | 2.27±0.07 | 3 |
| 27 | A21B12 | 2.27±0.05 | 4 |
| 28 | A13B21 | 2.26±0.07 | 3 |
| 29 | A10B10 | 2.25±0.06 | 3 |
| 30 | A20B21 | 2.25±0.03 | 3 |
| 31 | A14B6 | 2.24±0.10 | 4 |
| 32 | A4B9 | 2.20±0.04 | 3 |
| 33 | A13B12 | 2.19±0.03 | 4 |
| 34 | A19B12 | 2.16±0.06 | 4 |
| 35 | A12B3 | 2.16±0.15 | 4 |
| 36 | A16B10 | 2.16±0.04 | 3 |
| 37 | A13B3 | 2.16±0.12 | 3 |
| 38 | A11B12 | 2.14±0.07 | 4 |
| 39 | A7B2 | 2.14±0.13 | 4 |
| 40 | A4B12 | 2.11±0.05 | 4 |
| 41 | A19B3 | 2.09±0.10 | 4 |
| 42 | A12B10 | 2.07±0.07 | 4 |
| 43 | A14B14 | 2.02±0.12 | 3 |
| 44 | A21B2 | 2.00±0.10 | 3 |
| 45 | A13B2 | 2.00±0.05 | 4 |
| 46 | A20B2 | 1.97±0.05 | 4 |
| 47 | A9B8 | 1.96±0.07 | 3 |
| 48 | A16B5 | 1.96±0.08 | 4 |
| 49 | A14B10 | 1.94±0.05 | 4 |
| 50 | A10B3 | 1.94±0.06 | 4 |
| 51 | A8B8 | 1.94±0.04 | 4 |
| 52 | A14B5 | 1.93±0.05 | 4 |
| 53 | A15B19 | 1.93±0.10 | 4 |
| 54 | A4B2 | 1.93±0.06 | 4 |
| 55 | A14B2 | 1.92±0.12 | 4 |
| 56 | A4B20 | 1.92±0.05 | 3 |
| 57 | A6B10 | 1.87±0.05 | 4 |
| 58 | A7B10 | 1.84±0.09 | 4 |
| 59 | A21B10 | 1.81±0.08 | 4 |
| 60 | A21B11 | 1.75±0.03 | 4 |
| 61 | A9B10 | 1.60±0.02 | 4 |

Bold letters represented the strains selected for fruiting body production.

### Fruiting body production from crossbred strains

The parental and 18 crossbred strains were cultured, and fresh weight, and mycelial colonization time were compared between the strains (Table 4). Among the 18 crossbred strains, A20B3, A4B6, A16B3, and A16B6 showed abnormal fruiting bodies, and A5B1, A20B12, A6B6, and A10B12 did not form fruiting bodies at all. The weight of the fresh fruiting body of parental strain HS5 in one bag was 134.57 ± 5.45g, which was 43.14 g more than that of HS4 (Table 4). The BEs of HS5 and HS4 were 40.78 ± 1.65% and 27.71 ± 1.59%, respectively, in the first flush (Table 4). Two crossbred strains (A10B4 and A14B4) were more productive than parental strain HS5 (Fig 2), and five crossbred strains were more productive than parental strain HS4. The most productive strain was A10B4, but the difference in productivity between A10B4 and A14B4 was not significant. The mycelial colonization time of HS5 was 30.82 ± 0.98days, which was 6.1 days longer than HS4 (Table 4). Among the crossbred strains, A10B5, A10B4, and A1B18 displayed the shortest mycelial colonization times at 24.36 ± 1.03, 25.18 ± 1.33, and 25.00 ±1.18 days, respectively. Comprehensive comparisons of yield, mycelial colonization time indicated that A10B4 possessed the most desirable traits of all of the crossbred strains.

**Table 4.**
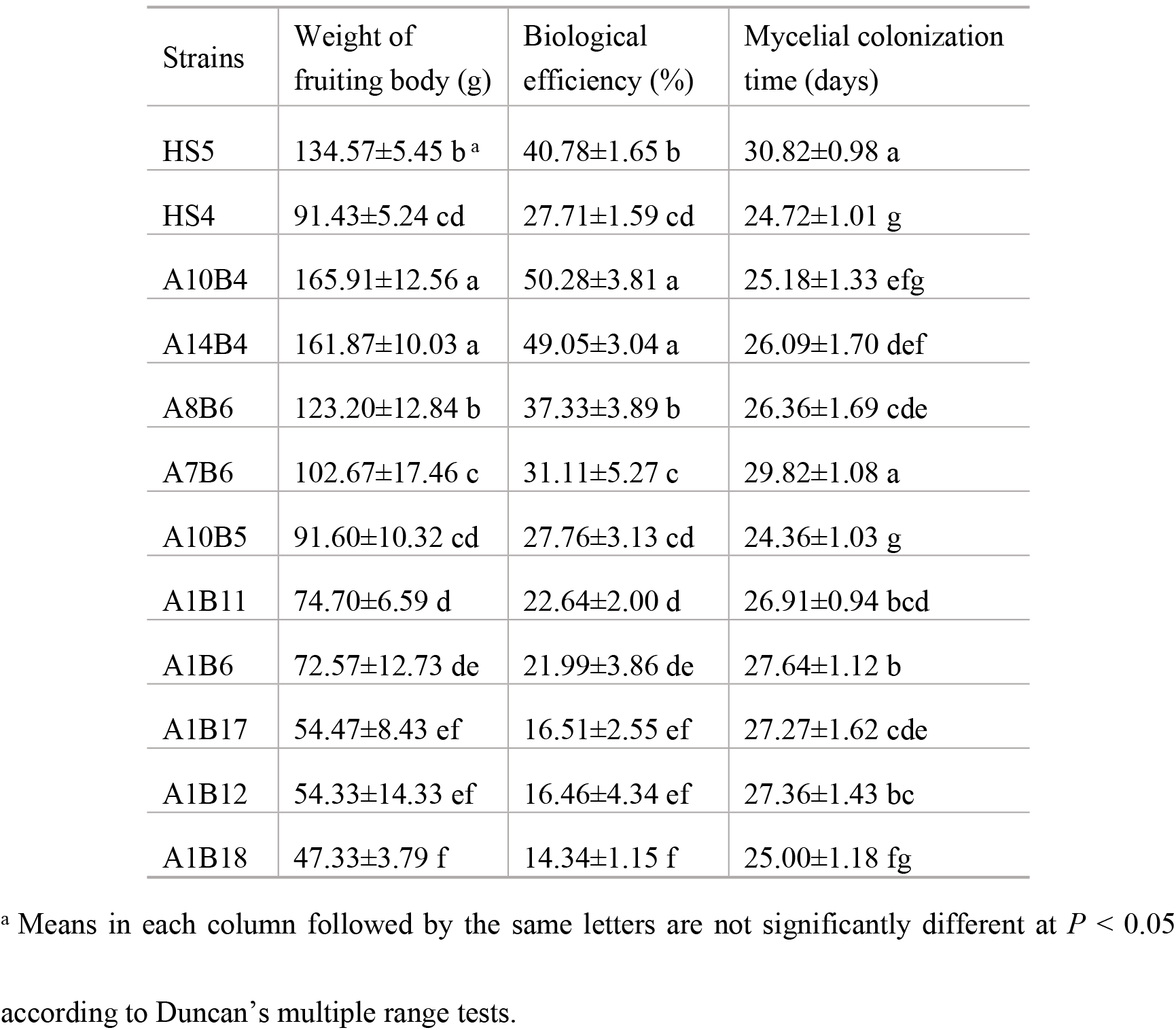
Comparison of the production yields and mycelial colonization times of fruiting bodies.

**Figure 2.**
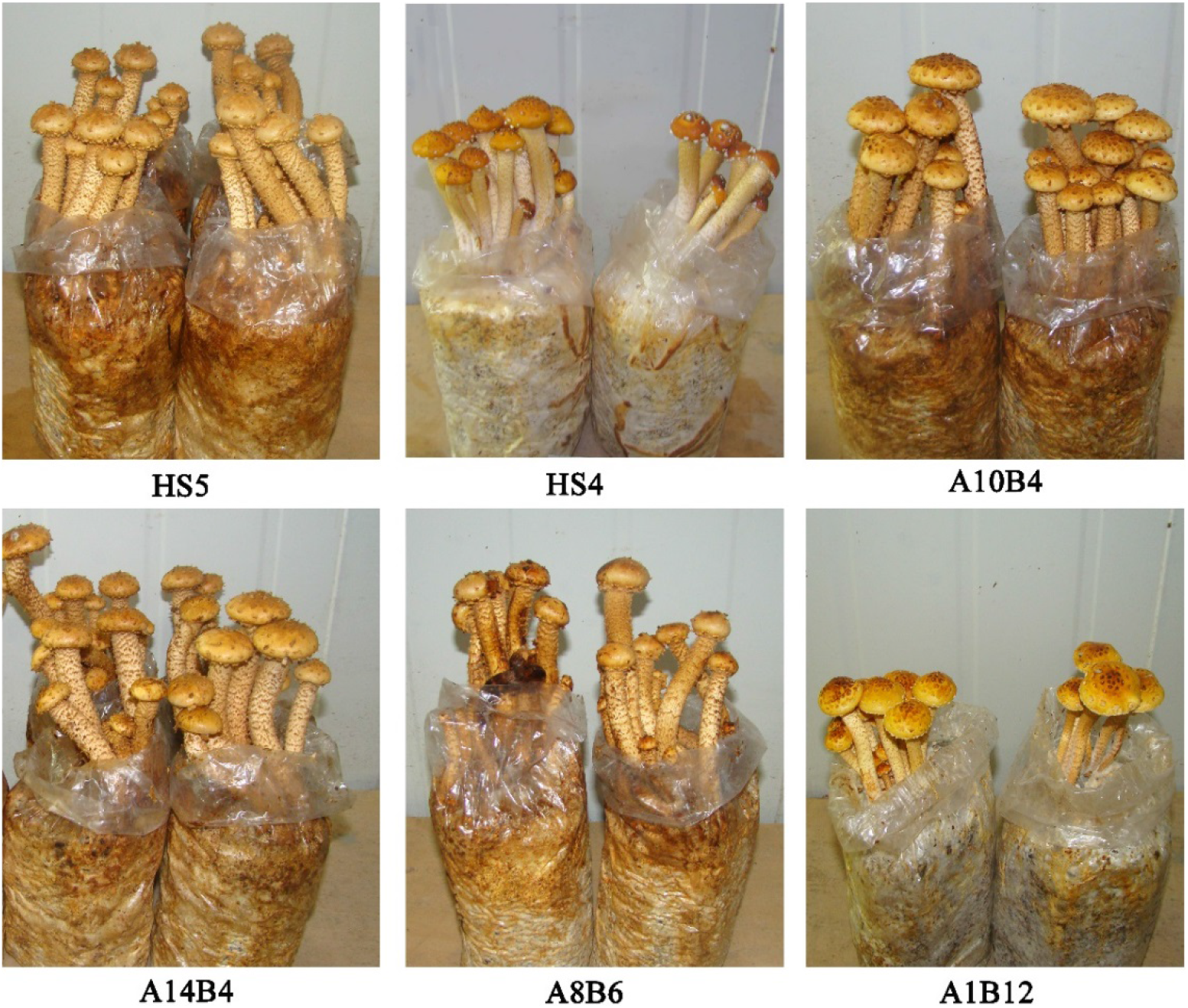
Morphological characteristics of HS4, HS5, and crossbred strain fruiting bodies.

### ISSR analysis

Crossbred strain A10B4 and the parental strains were analyzed by ISSR. Among the tested primers, four primers P11, P856, P23, and P857 were found to efficiently amplify the genomic DNA from all strains (Fig 3). The size of the polymorphic fragments obtained ranged from 200 bp to 5000 bp, and differences were observed in the number of bands obtained (Fig 3). The crossbred strain and two parental strains showed characteristic differences in the presence and absence of fragments. More than eight different fragments were amplified in crossbred strain A10B4 using the four primers. The results of PCR amplification indicated that A10B4 is a new hybrid strain and that differences in numbers of fragments are useful to identifying this strain.

**Figure 3.**
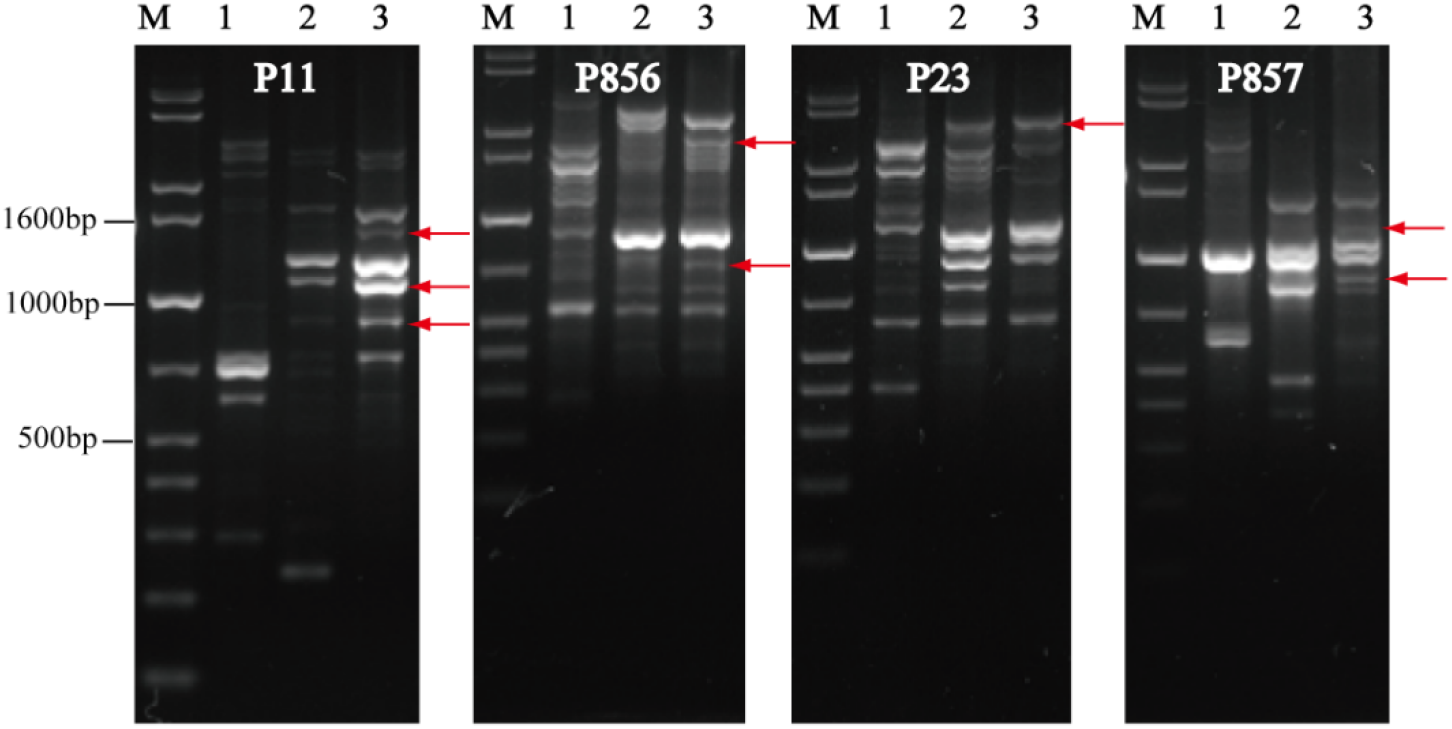
Amplification products obtained by PCR using ISSR primers and DNA from HS4, HS5, and A10B4. M, marker; line 1, HS4; line 2, HS5; line 3, A10B4. The arrows indicate the differenent fragments.

## Discussion

Cross-breeding using monokaryons was an efficient method to develop new strains. In this study, *P. adiposa* and *P. limonella* were crossbred to develop a new strain (*P. adiposa-limonella*) with high yield and MGR. The new high-yield crossbred strains obtained in this study should benefit future commercial cultivation efforts.

Dikaryotic mycelia form a clamp connection [12, 22], and in this study, mycelia were screened by detecting clamp connections under a microscope to identify dikaryotic hybrids. White rot fungi have an array of extracellular ligninolytic enzymes that synergistically and efficiently degrade lignin [21]. RBBR is an anthra-quinone-based dye with a structural resemblance to some polycyclic aromatic hydrocarbons [19]. Ligninolytic enzymes can decolorize RBBR. Sun et al. (2014) reported that ligninolytic enzymes can be used as a screening marker for new species of edible fungi, resulting in a screening method that is simple and direct. Selecting crossbred strains is an important step in breeding fungi, and, to reduce work and increase efficiency, we used RBBR fading degree (representing ligninolytic enzyme activity) and MGR to screen new strains in this study. One hundred and twenty-six and 103 monokaryons were isolated from *P. adiposa* HS5 and *P. limonella* HS4, respectively, and these monokaryons were selected based on ligninolytic enzyme activity and MGR, thus reducing breeding time compared with that needed only with clamp connection. The parental strain *P. adiposa* HS5 showed a higher yield and greater ligninolytic enzyme activity than *P. limonella* HS4. Six hybrid strains with high MGRs but relatively low ligninolytic enzyme activities (fading degrees ≤ 2) were also selected for mushroom production, and strains A1B18, A1B17, A1B11, A1B12, and A1B6 displayed low BEs, in general, with ligninolytic enzyme activities positively correlated with yield. The morphological characteristics of crossbred strains showed that monokaryon A1 was more closely related to HS4 than HS5. Among the 18 crossbred strains, four strains produced abnormal fruiting bodies, and four strains did not form fruiting bodies at all. The reason may be that *P. adiposa* and *P. limonella* are different species, and some of the resulting hybrids strains were infertile. To our knowledge, this is the first report of hybridization between *P. adiposa* and *P. limonella*. Ten crossbred strains formed normal fruiting bodies (Table 4). Strain A10B4 displayed the highest BE, as well as the shortest mycelial colonization time. Therefore, it was selected as the highest performing of the crossbred strains. ISSR and antagonistic activity assays confirmed that A10B4 is a new strain. Because the morphological characteristics of A10B4 are more like *P. adiposa* HS5, this strain was named *P. adiposa-limonella* HS54.

## Conclusion

In conclusion, a new crossbred strain, *P. adiposa-limonella* HS54, demonstrated favorable traits in terms of yield and BE. From a commercial point of view, the new strain has a clear advantage over the parental strains and suggests potential for future commercial cultivation of *Pholiota*.

## Acknowledgements

This work was supported by Beijing Agriculture Innovation Consortium (BAIC05-2019), the Open Research Fund Program of Beijing Key Lab of Plant Resource Research and Development, Beijing Technology and Business University (PRRD-2019-YB3).

## Author Contributions

**Conceptualization:** Shouxian Wang, Yu Liu.

**Investigation:** Chengbo Rong, Shuang Song, Li Yang, Yurong Niu, Xuejiao Pan.

**Formal analysis:** Chengbo Rong, Shuang Song, Li Yang, Qinggang Song.

**Methodology:** Jiachan Zhang, Li Yang.

**Software:** Jiachan Zhang, Chengbo Rong, Shuang Song.

**Supervision:** Shouxian Wang, Yu Liu.

**Writing – original draft:** Chengbo Rong, Shuang Song.

**Writing – review & editing:** Shouxian Wang, Yu Liu.

